# Complete characterization of the human immune cell transcriptome using accurate full-length cDNA sequencing

**DOI:** 10.1101/761437

**Authors:** Charles Cole, Ashley Byrne, Matthew Adams, Roger Volden, Christopher Vollmers

## Abstract

The human immune system relies on highly complex and diverse transcripts and the proteins they encode. These include transcripts for <u>H</u>uman <u>L</u>eukocyte <u>A</u>ntigen (HLA) class I and II receptors which are essential for self/non-self discrimination by the immune system as well as transcripts encoding B cell and T cell receptors (BCR and TCR) which recognize, bind, and help eliminate foreign antigens.

HLA genes are highly diverse within the human population with each individual possessing two of thousands of different alleles in each of the 9 major HLA genes. Determining which combination of alleles an individual possesses for each HLA gene (high-resolution HLA-typing) is essential to establish donor-recipient compatibility in organ and bone-marrow transplantations. BCR and TCR genes in turn are generated by recombining a diverse set of gene segments on the DNA level in each maturing B and T cell, respectively. This process generates <u>a</u>daptive <u>i</u>mmune <u>r</u>eceptor <u>r</u>epertoires (AIRR) of composed of unique transcripts expressed by each B and T cells. These repertoires carry a vast amount of health relevant information. Both short-read RNA-seq based HLA-typing^1^ and adaptive immune receptor repertoire sequencing^2–5^ currently rely heavily on our incomplete knowledge of the genetic diversity at HLA^6^ and BCR/TCR loci^7,8^.

Here we used our nanopore sequencing based <u>R</u>olling Circle <u>to</u> <u>C</u>oncatemeric <u>C</u>onsensus (R2C2) protocol^9^ to generate over 10,000,000 full-length cDNA sequences at a median accuracy of 97.9%. We used this dataset to demonstrate that deep and accurate full-length cDNA sequencing can - in addition to providing isoform-level transcriptome analysis for over 9,000 loci - be used to generate accurate sequences of HLA alleles for HLA allele typing and discovery as well as detailed AIRR data for the analysis of the adaptive immune system without requiring specific knowledge of the diversity at HLA and BCR/TCR loci.

## INTRODUCTION

The human immune system relies on highly diverse and complex receptors to protect us from a wide array of pathogens. The transcripts encoding these immune receptors are of great interest to basic and translational research as well as diagnostic and other clinical purposes^5,11,12^. However, RNA-seq, the current gold-standard for whole transcriptome analysis, fails short of describing these immune receptors completely and accurately^7^. Accurate and deep full-length cDNA sequencing of immune cell transcriptomes could overcome this shortfall by providing 1) the isoforms of surface receptors targeted in immunotherapy, 2) allele-resolved HLA transcript sequences central to self/non-self discrimination, and 3) B cell receptor (BCR) and T cell receptor (TCR) repertoires instrumental to the adaptive immune response to pathogens.

First, full-length cDNA sequencing should be capable of investigating the transcript isoforms of surface receptors expressed by B cells which have important roles in the immune response but are also themselves targets in the treatment of B cell derived leukemia. For example, current antibody and <u>C</u>himeric <u>A</u>ntigen <u>R</u>eceptor (CAR) T-cell therapies against B-cell acute lymphoblastic leukemia (B-ALL) target epitopes of CD19, CD20, and CD22^13,14^. However, evidence is accumulating that these epitopes might be absent in some transcript isoforms of these genes. Detailed analysis of this absence and possible consequences is hampered by the inability of RNA-seq to resolve transcript isoforms^10,15,16^. Determining isoform level transcriptomes of healthy and cancerous immune cells might therefore inform treatment decisions and future development.

Second, full-length cDNA sequencing could be used to accurately determine the sequence and identity of alleles of HLA genes. HLA genes encode for the <u>M</u>ajor <u>H</u>istocompatibility <u>C</u>omplex (MHC) group of immune receptors which are instrumental in the presentation of antigens on the cell surface. Most cells express HLA class I transcripts (A, B, and C) which encode for proteins that present fragments of other endogenous proteins and can therefore alert the adaptive immune system of viral infections hijacking the cellular machinery. Professional antigen presenting cells like B cells, dendritic cells, and Macrophages also express HLA-class II (including DPA1, DPB1, DRA, DRB1, DQA1, and DQB1) transcripts which encode proteins that present foreign antigens on the cell surface and can lead to the activation of the adaptive immune system^17^. Determining the identities of HLA alleles that are present within an individual’s genome (HLA-typing) is of central importance to establish compatibility of donor and recipient for organ and bone-marrow transplantation because the genes encoding HLA transcripts are highly diverse throughout the human population and will therefore be recognized as foreign if not properly matched.

Currently, HLA-typing is performed in clinical laboratories by amplifying the genomic DNA encoding these genes and sequencing these amplicons with short read sequencers^1^. This is required because even though all of these genes are expressed by immune cells in the blood and are therefore captured by RNA-seq, even sophisticated computational tools relying on complex workflows and statistically rigorous frameworks struggle to process RNA-seq data and provide reliable allelic identities^6,18^. Further, these tools rely on databases of known HLA allele sequences which prevents them for identifying so far unknown HLA alleles. Extracting accurate and full-length HLA allele sequences from full-length cDNA sequences could therefore dramatically simplify the computational task of determining the identities of HLA alleles present in a sample, independent of whether these HLA alleles have previously been identified.

Third, accurate full-length cDNA sequencing should also be capable of determining adaptive immune receptor sequences, including BCR (antibody) heavy and light chains and TCR alpha and beta transcripts. Due to their complexity, these transcripts likely pose the biggest challenge in transcriptome analysis. These receptor transcripts contain a constant and a variable region. The constant region determines the type and characteristics of the receptor and the variable region determines its binding affinity. The exon encoding these variable regions is generated through the process of somatic VDJ recombination which randomly recombines of a pool of similar but distinct V(, D), and J gene segments^19^. Each developing T or B cell uses this somatic recombination process unique to these cell types to rearrange non-functional loci into two functional genes (B cells: heavy/light (kappa of lambda), T cells: alpha/beta). Each T and B cell thereby generates a unique TCR and BCR/Antibody. The repertoires of these transcripts present in blood or tissue samples carry a large amount of information on the composition of the loci they are expressed from, as well as the activation state, clonal composition (including malignant clones in Leukemia)^5^, and basic biological processes of the adaptive immune system.

Currently, targeted <u>A</u>daptive Immune <u>R</u>eceptor <u>R</u>epertoire <u>seq</u>uencing assays (AIRR-seq) methods are routinely employed to sequence these transcripts and investigate the human immune system^20–22^. Specialized assays are required because the diversity and unique nature of these transcripts make them practically impossible to analyze at full length using standard RNA-seq protocols. Further, the majority of these assays are based on primers against known V segments and therefore potentially biased against so far unknow V segments. The ability to instead extract these full-length transcripts in an unbiased way from whole transcriptome full-length cDNA sequencing would greatly simplify workflows and expand the information that can be recovered from non-targeted transcriptome analysis.

Here we show that our previously published R2C2 method^9,23^ implemented on the Oxford Nanopore Technologies MinION sequencer can not only provide high quality isoform-level whole-transcriptome analysis but also provide allele-resolved HLA sequences suitable for high resolution HLA-typing and replace specialized AIRR-seq methods for many applications. To this end, we analyzed RNA extracted from a human <u>P</u>eripheral <u>B</u>lood <u>M</u>ononuclear <u>C</u>ells (PBMC) sample – a mix of mostly monocytes, B cells, and T cells. We generated 10,000,000 R2C2 reads that covered cDNA molecules end-to-end. We analyzed these reads using our Mandalorion tool to identify transcript isoforms and implemented new tools to determine allele-resolved sequences of full-length HLA transcripts. We also used a new computational pipeline to extract BCR and TCR sequences from these reads which we processed to generate AIRR data.

Our results show that accurate and deep full-length cDNA can resolve the most complex transcripts in the mammalian genome and represents an unbiased alternative to current HLA-typing and AIRR-seq methods.

## RESULTS

We extracted total RNA and genomic DNA from PBMC samples purified from the whole blood of a healthy male adult. DNA was used for high-resolution HLA-typing whereas total RNA was used for several transcriptome analysis assays. First, we generated full-length cDNA using a modified first half of the Smart-seq2 protocol^24^. cDNA was then split to generate sequencing libraries with three different methods. First, to generate standard Smart-seq2 libraries, part of this cDNA was then tagmented using Tn5^25^. Next, we circularized the full-length cDNA and performed rolling circle amplification on the resulting circular DNA. This reaction generated long dsDNA containing multiple concatemeric copies of the original full-length cDNA. We either sequenced this DNA on the ONT MinION using the R2C2 protocol^9^ or tagmented it with Tn5 to generate a hybrid Smart-seq2 and R2C2 short-read library we named Smar2C2 (Fig.1).

**Fig. 1:**
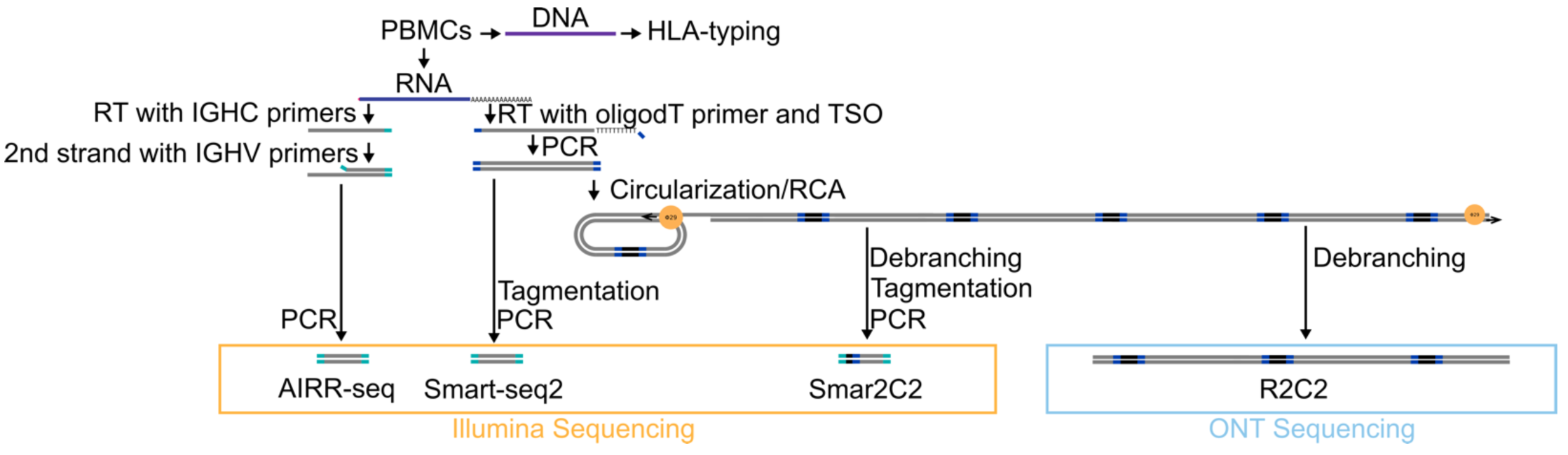
Analysis of the adaptive immune system through high-throughput sequencing. Schematic of experiment design. DNA and RNA extracted from a PBMC sample underwent several library preparation protocol to generate AIRR-seq, Smart-seq2, Smar2C2, and R2C2 libraries that were sequenced on Illumina and ONT sequencers.

### R2C2 data characteristics

The R2C2 method sequences the same cDNA sequences multiple times to overcome the low accuracy of raw 1D ONT cDNA reads^26^. It does so by circularizing double-stranded cDNA molecules using a DNA splint in a variant of Gibson assembly^27^ (NEBuilder, New England Biolabs) and amplifying this circularized cDNA using phi29-based rolling circle amplification (RCA). The resulting RCA product consists of concatemers of the same cDNA sequence that can be sequenced on long-read sequencers and converted into accurate consensus sequences. In an update to the previously published version^9^ of R2C2 we also incorporated the use of unique molecular identifiers (UMIs) as part of the DNA splints. Because the UMIs are linked to cDNA after PCR amplification they do not indicate unique RNA molecules, but instead reflect unique circularization events, thereby allowing us to combine the raw reads originating from the same cDNA molecule to improve R2C2 consensus read accuracy.

We generated R2C2 data in four technical replicates using individual ONT MinION 9.4.1 flow cells. 1D raw reads from each flow cell were processed using C3POa tools^9^ we developed previously to generate a total of 10,298,086 R2C2 consensus reads. Of these reads, 122,353 could be grouped with one or more other reads based on their UMIs and were combined into 58,893 R2C2-UMI reads. We aligned R2C2 and R2C2-UMI consensus reads to the human genome (hg38, primary assembly) using *minimap2*^28,29^. Analysis of the resulting alignments determined an accuracy (Matches/Matches+Mismatches+Indels) of 97.9% for regular R2C2 reads and of 99.3% for R2C2-UMI reads. Because of the relatively low number of these R2C2-UMI reads, we merged regular R2C2 reads and R2C2-UMI reads into a single data set with a total number of 10,234,626 reads of which >99.9% aligned to the human genome.

We used featureCounts^30^ to quantify gene-level expression based on these R2C2 reads and Smart-seq2 reads. We found that overall the different protocols correlated well for the analysis of the same cDNA sample with a Pearson r-value of 0.93. While this shows that R2C2 captures the transcriptome in a quantitative manner it is not the focus of this analysis as the strength of long-read sequencing like R2C2 is not gene-level but instead isoform-level transcriptome analysis.

### Isoform identification and evaluation

To identify transcript isoforms, we used the R2C2 reads as input into a revised version of the Mandalorion tool (v3, github)^10^. We detected 21,358 transcript isoforms expressed from 9,971 gene loci. The isoform sequences Mandalorion produced showed a median accuracy of 99.7%. Their actual accuracy is likely even higher considering the genome sequence of the sample donor is not expected to be identical to the human reference genome sequence.

We further evaluated the quality of these isoforms using SQANTI (sqanti_qc.py)^31^. 12,250 isoforms were scored as full-splice matches of known transcripts although with potentially new transcription start and polyA sites. 2,265 were scored as novel-in-catalog, meaning that they used known splice-sites in previously un-annotated combinations. 1,369 were scored as novel-not-in-catalog, which means that they contain un- annotated splice sites and potentially entire exons. 4,661 were scored as incomplete-splice-match which could mean that they are potential artifacts and were not used to extract potentially new TSS and polyA sites. Overall, we detected 5,525 new TSSs, 5,712 new polyA sites, and 365 new exons not present in the gencode(v29) annotation^32^.

We then tested whether we could validate these new features using short-read Illumina protocols. Because all general-purpose RNA-seq protocol struggle to capture transcript ends, we developed a new Illumina-based RNA-seq method that overcomes this issue. This Smar2C2 method is a hybrid of the <u>Smar</u>t-seq2 and R<u>2C2</u> methods that tagments not cDNA molecules but the cDNA concatemers generated as part of the R2C2 method. As a result, Tn5 based tagmentation is not affected by cDNA ends because these ends are now encapsulated within a much larger DNA molecule. We found that the Smar2C2 protocol does not distort the cDNA composition, as data produced from Smar2C2 correlates very well with Smart-seq2 data for gene-level expression analysis with a Pearson r-value of 0.97 (Fig. 1B). Further, processing of the ∼39 million Smart-seq2 and ∼23 million Smar2C2 read pairs suggests that Smar2C2 data contains ∼6x (Smart-seq2: 3% and Smar2C2 17% of all reads) more reads covering a putative TSS (pTSS) and ∼25x (Smart-seq2: 0.17% and Smar2C2 4% of all reads) more reads covering a putative polyA (pPolyA) site than standard Smart-seq2 data. Smar2C2 read coverage dropped sharply outside new splice sites, and once extracted, pTSS and pPolyA read coverage also dropped sharply outside new TSS and polyA sites, thereby validating the new features we identified as present within the cDNA pool (Fig. 2C).

**Fig. 2:**
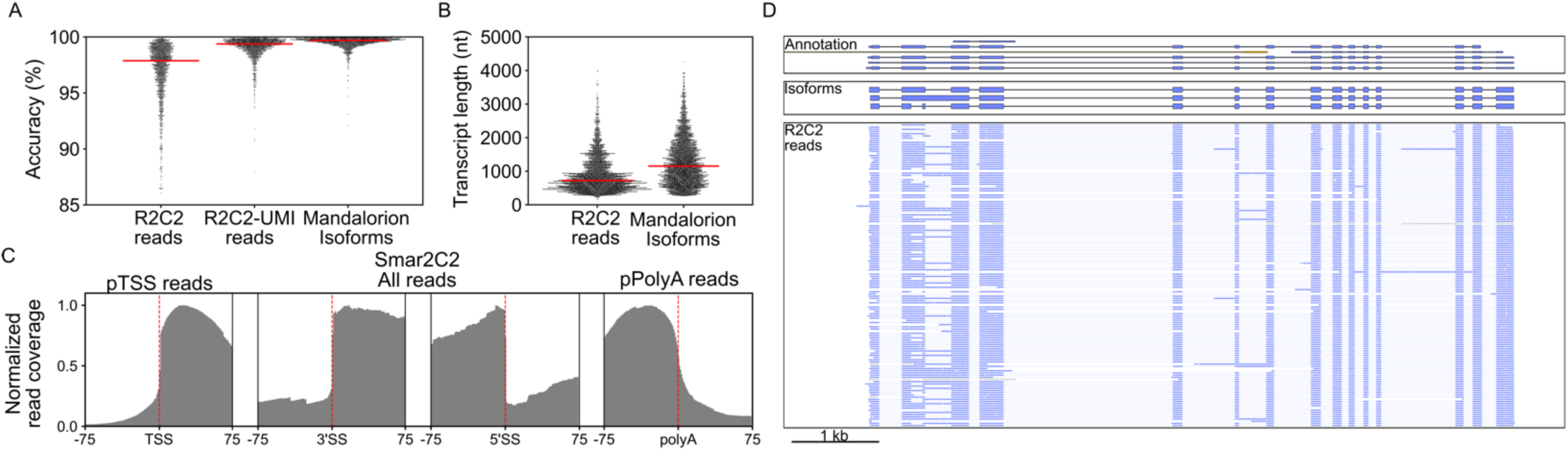
Isoform level transcriptome analysis. Swarmplots of the A) accuracy [*Matches/(Matches+Mismatches+Indels)*] and B) transcript length [*Matches+Mismatches*] of R2C2 reads (with and without UMI) and isoforms. Red line indicates median. C) Normalized read coverage around newly identified splice sites, Transcription Start Sites (TSSs), and polyA sites determined by either all (splice sites), pTSS (Transcription start sites) or pPolyA (PolyA sites) Smar2C2 reads. D) The CD19 gene locus is shown in a genome-browser view. Gencode annotation (top), Mandalorion isoforms (center) and R2C2 reads (bottom) are shown. Direction of shown features is indicated by color (“+” => blue, “-” => yellow).

To showcase the usefulness of deep full-length isoform data we highlight the CD19 gene expressed by B cells. CD19 is of great importance because it is a target of cancer immunotherapy in B cell <u>A</u>cute <u>L</u>ymphoblastic <u>L</u>eukemia (ALL) and other B cell cancers. At a total read depth of ∼10 million reads, only 146 reads aligned to the CD19 gene. These 146 R2C2 reads appeared to split into 3 major and several minor isoforms. Mandalorion identified the 3 major isoforms which confirmed a previously identified splice site in the 2nd exon of CD19 that we recently observed in single B cells^9^. Because they encode distinct proteins, these isoforms may affect whether CAR T-cells^15,16^ can bind leukemia cells expressing them.

### Allele specific isoform expression

At close to 98% accuracy, R2C2 reads should be well suited for allele-specific isoform expression analysis. Because the SNP identification is much more established with short-read data, we made use of the Smart-seq2 and Smar2C2 data that we produced to identify SNPs present in the PBMC sample we analyzed using the standard <u>G</u>enome <u>A</u>nalysis <u>T</u>ool<u>k</u>it (GATK) RNAseq workflow^33^. Using the TurboPhaser.py script we developed for this purpose, we then extracted heterozygous SNPs from this list and phased these SNPs within gene boundaries using R2C2 reads. TurboPhaser.py then sorted R2C2 reads and short-read RNA-seq read pairs into alleles based on the phased SNPs they contained. Overall, we assigned 756,072 R2C2 reads (7.4% of all reads, Allele1: 377,794; Allele2: 378,278) and 1,817,151 RNA-seq read pairs (2.9% of all read pairs, Allele1: 872,130; Allele2: 945,021) to either of two alleles. It is noteworthy that R2C2 reads are more than twice as likely to be sorted into alleles than RNA-seq read pairs based on the same set of phased SNPs. This is likely due to R2C2 reads, in contrast RNA-seq reads, covering entire transcripts and all SNPs they contain.

Next, we used the allele-resolved R2C2 reads to quantify the expression of the previously identified isoforms. To this end, we separated these reads into the four technical replicates based on the flow cells they were generated on. Using DESeq^34^, we then identified ∼80 isoforms that showed differential expression between the two alleles while accounting for the technical variation associated with each minion run. The SNHG5 gene highlights this differential expression with transcripts of one allele always retaining the first intron of the gene and transcripts of the other allele either splicing or retaining the intron (Fig. 3B). Interestingly, seven of the ∼80 differentially expressed isoforms originated from an HLA gene. HLA-A and HLA-DPA1 both show differentially expressed isoforms with alternative polyA sites. While the alternative HLA-A isoform was previously observed^35,36^, the HLA-DPA1 isoform was not (Fig. 3C,D).

**Fig. 3:**
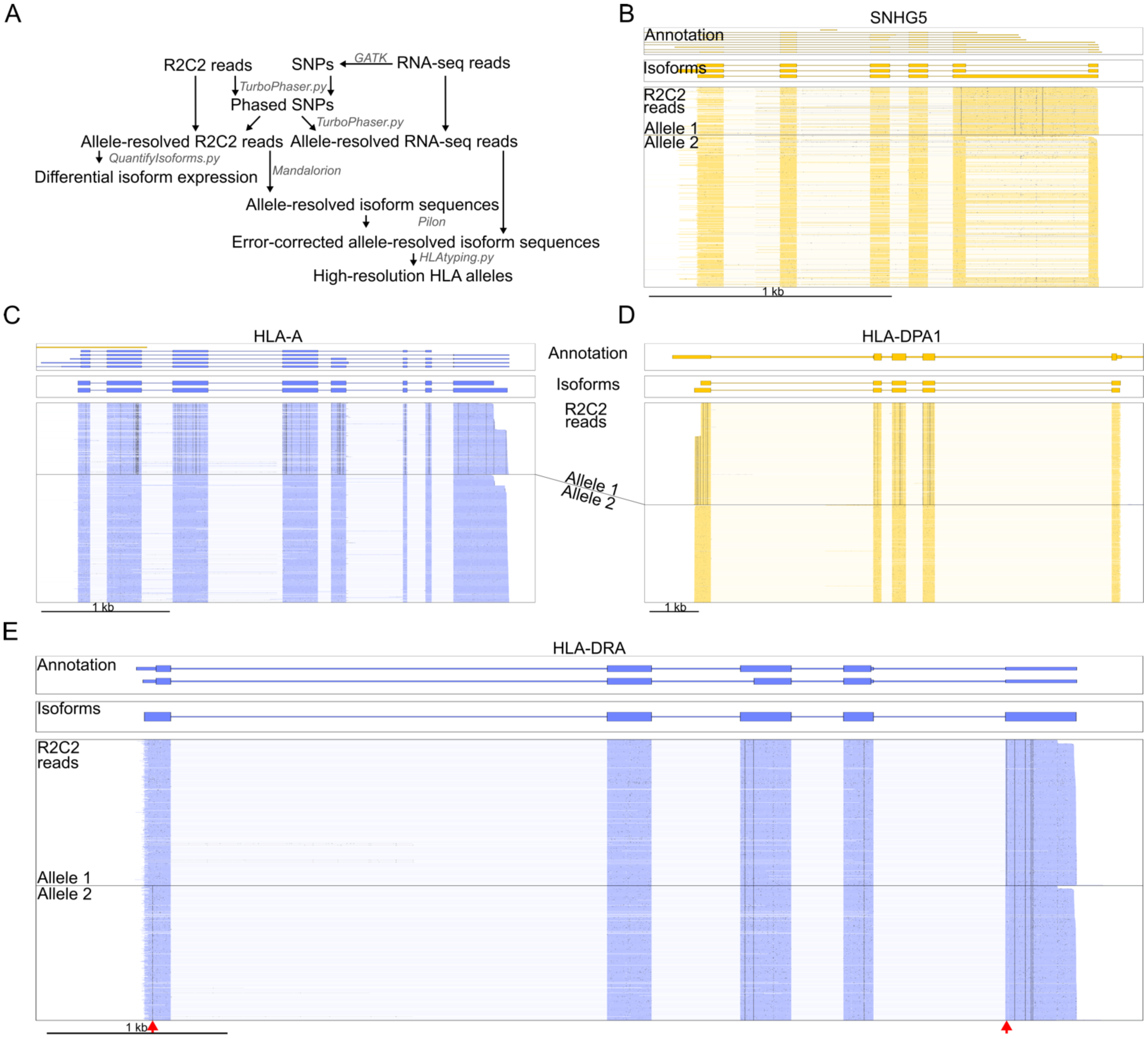
Allele-resolved isoform expression and sequences. A) Computational strategy for determining allele-resolved isoforms. B-E) SNHG5, HLA-A, HLA-DPA1, and HLA-DRA gene loci are shown in a genome-browser view. Gencode annotation (top), Mandalorion isoforms (center) and allele-resolved R2C2 reads (bottom) are shown. Direction of shown features is indicated by color (“+” => blue, “-” => yellow). Mismatches of R2C2 reads to the genome reference are shown in black. Both HLA-A and HLA-DPA show allele-resolved differential expression of isoforms with alternative polyA sites. E) Red arrows indicate allele-specific variants in the HLA-DRA gene.

### Allele-resolved isoform sequences enable high resolution HLA-typing

Next, we investigated whether allele-resolved R2C2 reads are suited for identifying which HLA alleles are present in an individual. HLA genes are highly diverse within the human population with different individuals carrying different alleles. Identifying which HLA alleles are present in an individual (HLA-typing) is of utmost importance when establishing donor-recipient compatibility in transplantation medicine. Current RNA-seq based HLA-typing methods rely on databases of previously identified and systematically cataloged HLA alleles. The IPT-IMGT/HLA database contains the systematic names and sequences of thousands of different HLA alleles. The systematic names (e.g. HLA-A*01:01:01:01) contain multiple groups of digits separated by colons to denote the relationship between sequences. The first two digits of an HLA allele name identify the group the allele belongs to (2-digit resolution), the second two digits identify the protein sequence and are affected by non-synonymous variants (4-digit resolution), the third group of digits indicates synonymous variants in the coding region of the protein (6-digit resolution), and the fourth group of digits variants identifies variants outside the protein coding region (8-digit resolution).

Using this database, current RNA-seq based methods can determine the identity of HLA alleles present in an individual with 4 to 6-digit resolution. However, even the most advanced methods like arcasHLA^18^ have a 10% error-rate for the identification of some HLA genes and, importantly, cannot determine new HLA alleles absent from the database they use. Reliable HLA typing therefore still requires dedicated DNA-based approaches. These approaches PCR-amplify full-length HLA genes from genomic DNA and determine the sequence of the resulting amplicons. These sequences are then compared to the database of known HLA alleles to determine 6-digit HLA types.

R2C2 should enable a similar approach to targeted DNA-based approaches by determining the full-length sequences of all HLA transcripts present in the analyzed sample without knowledge of what HLA alleles have previously been identified. To test this, we used the 377,794 R2C2 reads assigned to the first allele and 378,278 R2C2 reads assigned to the second allele as separate inputs into Mandalorion which then generated 2,237 and 2056 allele-specific isoforms, respectively. Importantly, Mandalorion generates entirely read-based consensus sequences for each isoform it identifies, which in this case included at least one full-length isoform for each major HLA gene on either allele. Next, to achieve the highest possible accuracy required for identifying variants and achieving unambiguous HLA-typing, we used allele-resolved RNA-seq reads to error-correct the allele-specific Mandalorion isoforms with pilon.

To determine the identity of the HLA alleles present in the analyzed sample, we used the HLAtyping.py script which aligns these error-corrected allele-resolved HLA isoform sequences to the complete database of HLA alleles^37,38^ using minimap2 and extracts the best match for each HLA gene. After finding the best HLA allele match of an error-corrected allele-specific isoform, we truncated the match to 6-digit resolution to match clinical high-resolution HLA-typing. All HLA alleles we identified in this way (R2C2+RNA-seq) matched DNA-based high-resolution HLA typing performed at the Immunogenetics and Transplantation Laboratory (ITL) at UCSF (Targeted Amplicon NGS) (Table 1).

**Table 1:** R2C2 full-length cDNA sequencing enables high-resolution HLA-typing. HLA alleles were typed using the programs indicated on top. Different programs employed for HLA-typing rely on different data sources. The HLA Twin program requires DNA-based amplicon sequencing (Targeted Amplicon NGS), Our HLA-typing.py program requires isoform sequences generated using fulllength cDNA (R2C2) and polished with RNA-seq sequences. The seq2HLA and arcasHLA program require only RNA-seq sequences. DRA was not evaluated by Targeted Amplicon NGS. Contradicting results are shown in red.

|  | Allele 1 |  |  |  | Allele 2 |  |  |  |
| --- | --- | --- | --- | --- | --- | --- | --- | --- |
| Data Source | Targeted Amplicon NGS | R2C2+RNA-seq | RNA-seq | RNA-seq | Targeted Amplicon NGS | R2C2+RNA-seq | RNA-seq | RNA-seq |
| Program | HLA Twin | HLAtyping.py | seq2HLA | arcasHLA | HLA Twin | HLAtyping.py | seq2HLA | arcasHLA |
| HLA-A | 03:01:01 | 03:01:01 | 03:01 | 03:01:01 | 32:01:01 | 32:01:01 | 32:01' | 32:01:01 |
| HLA-B | 35:01:01 | 35:01:01 | 35:01 | 35:01:01 | 39:01:01 | 39:01:01 | 39:01 | 39:01:01 |
| HLA-C | 04:01:01 | 04:01:01 | 04:01 | 04:01:01 | 12:03:01 | 12:03:01 | 12:03 | 12:03:01 |
| HLA-DRA |  | 01:01:01 | 01:01 | 01:01:01 |  | 01:01:01 | 01:01 |  |
| HLA-DRB1 | 16:01:01 | 16:01:01 | 16:01 | 16:01:01 | 01:01:01 | 01:01:01 | 01:01 | 01:01:01 |
| HLA-DPA1 | 01:03:01 | 01:03:01 | 02:01 | 01:03:01 | 02:01:01 | 02:01:01 | 02:01 | 02:01:01 |
| HLA-DPB1 | 04:02:01 | 04:02:01 | 105:01 | 04:02:01 | 10:01:01 | 10:01:01 | 10:01 | 10:01:01 |
| HLA-DQA1 | 01:01:01 | 01:01:01 | 01:02 | 01:01:01 | 01:02:02 | 01:02:02 | 01:02 | 01:02:02 |
| HLA-DQB1 | 05:01:01 | 05:01:01 | 05:01 | 05:01:01 | 05:02:01 | 05:02:01 | 05:01 | 05:02:01 |

Having confirmed the accuracy of the HLA alleles we identified, we compared them to HLA alleles determined using only RNA-seq data and different programs. The seq2HLA package (SS2/seq2HLA) only generates data with 4-digit resolution and failed to identify HLA-DPA1, HLA-DQA1 and HLA-DQB1 as heterozygous an miscalled HLA-DPB1. The arcasHLA^18^ package (RNA-seq/arcasHLA) performed better, determining the correct HLA alleles for all HLA genes. Interestingly however, while both RNA-seq/arcasHLA and R2C2+RNA-seq strategies identify only one 6-digit resolution allele for the HLA-DRA genes (HLA- DRA*01:01:01), only R2C2+RNA-seq identifies the HLA-DRA gene as heterozygous by identifying distinct sequences for the two alleles. Visualizing allele-resolved R2C2 reads shows heterozygous variants of these alleles that are several hundred bp apart. ArcasHLA cannot resolve this because the variants are outside the protein coding region, where they affect 8-digit but not 6-digit HLA resolution (Fig. 3E).

Overall, these findings show that accurate full-length cDNA sequencing at high depth allows the determination of highly accurate sequences of HLA alleles which can then be used for high-resolution HLA-typing. In contrast to short-read RNA-seq based HLA-typing which requires a reference database, these HLA allele sequences can be also used to discover so far unknown HLA alleles. Beyond clinical applications, full-length cDNA sequencing therefore represents a potentially valuable research tool for investigating the diversity of the human population which currently requires high coverage whole genome sequencing data.

### Extracting adaptive immune receptor repertoires (AIRR) from R2C2 data

Next, we evaluated whether accurate full-length sequencing could also - completely or in part - replace specialized assays for the analysis of adaptive immune receptor repertoires. To do so, we extracted R2C2 reads from our data set which aligned to BCR or TCR gene loci. Because each read could be derived from a unique AIRR transcript, every read has to be treated independently and we cannot rely on isoform determination. Further, since the loci present in the human genome sequence are not the rearranged loci we find in each individual B or T cell, we do not evaluate these alignments directly. Instead we took the sequences of these reads and annotated them using the IgBlast^39^ algorithm which identifies V(, D), and J segments, CDR3 sequences at the V(D)J intersection, and mutations present in each sequence. Finally, we use sequence similarity to determine which constant region is present in each sequence. In this way, we identified tens of thousands of adaptive immune receptor sequences (Table 2).

**Table 2:** AIRR data can be extracted from R2C2 reads. Number of reads that aligned to the respective locus and could be annotated as AIRR transcripts using IgBlast is shown.

| Receptor | Gene | Number of R2C2 reads |
| --- | --- | --- |
| BCR | IGH | 12,863 |
|  | IGL | 26,759 |
|  | IGK | 24,460 |
| TCR | TRA | 7,289 |
|  | TRB | 13,118 |
|  | TRD | 316 |
|  | TRG | 551 |

We then performed in-depth analysis on these annotations with a focus on BCR heavy chain (IGH) sequences which are the only adaptive immune receptor undergoing VDJ recombination, somatic hypermutation, and class-switch recombination^19^. This makes BCR IGH sequences the biggest analysis challenge and therefore the best and most conservative target for detailed validation.

To evaluate how well this R2C2-based approach performed, we compared the resulting R2C2-based BCR IGH repertoires to 266,390 BCR IGH sequences we generated from the same RNA sample by the gold-standard UMI-based targeted AIRR-seq method we developed (called AIRR-seq from here on out)^3,12,20,21,40^. We also used 3,261 BCR IGH sequences generated from a different RNA sample of the same individual that we published previously^2^. These sequences were generated using our TMI-seq method which, in contrast to standard AIRR-seq^3^, succeeds in covering the entire V segment by overcoming Illumina read length limitations through a combination of Tn5-based tagmentation and unique molecular identifiers. We focused our analysis on the most relevant features of the BCR repertoire, namely 1) CDR3 length, 2) Isotype usage, 3) V segment composition and usage, 4) Somatic Hypermutation, and 5) Clonality.

#### CDR3 length

First, we investigated whether R2C2-based BCR IGH repertoires capture the full width of CDR3 lengths. The sequences of CDR3s are responsible for the majority of an BCR/antibodies specificity and are composed of semi-random sequence at the intersection of V, D, and J segments. However, CDR3 sequences in functional antibodies are limited to certain lengths to maintain the reading frame between variable and constant regions. This limitation can be clearly observed in CDR3 lengths of AIRR-seq sequences but is less pronounced for R2C2 sequences. This is likely due to remaining Indel errors in the CDR3 sequences (Fig 4A). So, although R2C2- based repertoires capture CDR3 length of the appropriate mean length and distribution, downstream analyses that rely on CDR3 length to differentiate functional and non-functional BCR sequence would be hampered.

**Fig. 4:**
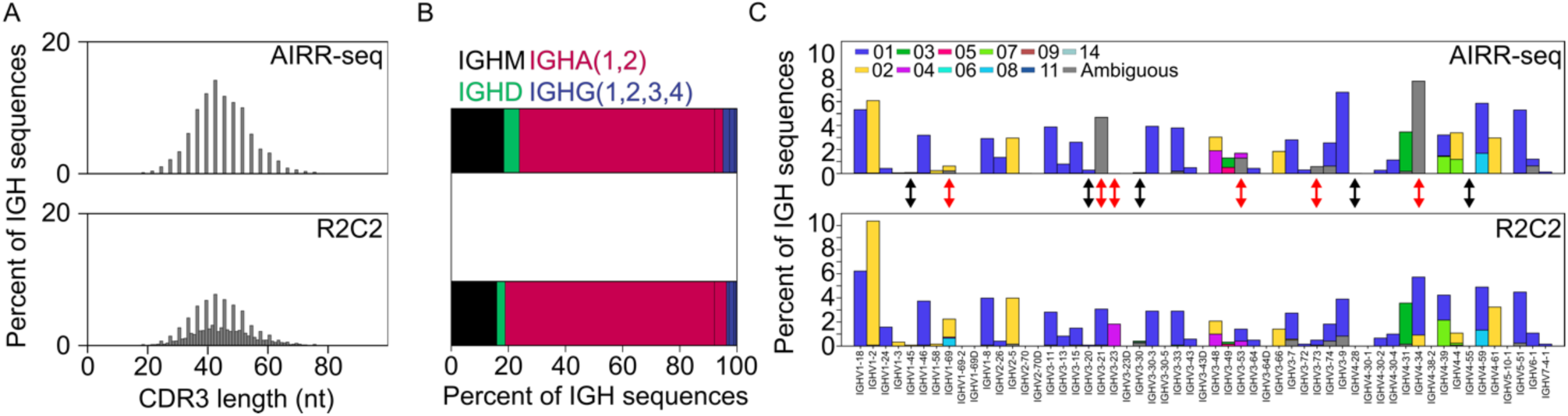
R2C2 repertoires have advantages and disadvantages compared to AIRR-seq repertoires: A) CDR3 lengths, B) Isotype distribution, and C) V segment usage are shown for BCR IGH repertoires are determined by AIRR-seq (top) and R2C2 (bottom). In C), different bar colors indicate different V segment alleles with grey indicating ambiguous allele calls. Red arrows indicate V segments where AIRR-seq fails to identify an allele unambiguously but R2C2 succeeds. Black errors indicate rarely recombined V segments that AIRR-seq but not R2C2 detected.

#### Isotype usage

Second, we investigated whether R2C2-based BCR IGH repertoires can be used to determine B cell isotype usage. Isotype usage reflects the activity of the adaptive immune system at a given time. IGHM and IGHD sequences are mostly expressed by naive B cells, while IGHA, IGHG and IGHE sequences are only expressed by previously activated B cells that have undergone class-switch recombination. Relative isotype expression has been shown to change upon vaccine administration^3^, infection, or immunosuppression^12^.

Isotype usage was similar between AIRR-seq and R2C2-derived repertoires suggesting that R2C2- derived repertoires will be able to determine the immune activation state of an individual faithfully (Fig. 4B). Improving upon any AIRR sequencing approach currently available, R2C2-derived repertoires also resolve whether a BCR IGH transcript encodes for a membrane-bound or secreted (antibody) protein. This showed that the majority of IGHM transcript are membrane-bound (Membrane (M): 1261; Secreted (S): 417) while, surprisingly, IGHD transcripts were split evenly between the two isoforms (M: 123; S:164). As expected, over 95% of the sequences of IGHA and IGHG isotype subtypes were secreted (e.g. IGHA1: M:182; S:8113).

#### V segment composition and usage

Third, we determined whether R2C2 repertoires could be used to investigate V segment usage. The adaptive immune receptor genes loci (BCR IGH, IGL, and IGK and TCR TRA, TRB, TRD, and TRG) are among the most diverse and complex in the human genome. Each BCR IGH locus is thought to contain 40-50 V segments and individuals diverge in which V segments and V segment alleles they possess. AIRR data can determine which V segments are present in an individual’s genome and how often they are recombined. However, standard AIRR-seq assays most often use PCR primers within V segments, thereby masking variation underneath the priming site and missing variation beyond it. TMI-seq addressed this by priming in the Leader exon beyond the V segment exon, however it still relied on potentially biased multiplex primers and requires complex molecular biology and computational workflows.

Because the R2C2-based repertoires could not be successfully analyzed with computational tools meant for virtually error-free AIRR-seq^41,42^, we determined V segment composition in the analyzed individual by simply counting how many reads were scored by IgBlast as using a specific V segment with three or fewer mismatches. Reads that assigned equally well to different alleles of the same V segment were counted as ambiguous while reads aligned equally well to different V segments were discarded for this analysis. A V segment allele was counted as detected in a repertoire if it was seen in at least two sequences and accounted for at least 20% of sequences of the V segment

In general, AIRR-seq and R2C2 showed similar recombination frequencies for V segments. However, we found that the deeper AIRR-seq data has an advantage when detecting V segment alleles for V segments that are rarely recombined, including IGHV1-45 (0.08% of IGH sequences in AIRR-seq data), IGHV3-20 (0.3%), IGHV3-30 (0.04%), IGHV4-28 (0.01%), IGHV4-55 (0.02%) which R2C2 did not detect at our requisite abundance of three independent reads. However, we found that R2C2 can unambiguously detect V segments that AIRR-seq could not (Fig 4C). AIRR-seq could not detect one or two of the alleles for IGHV1-69, IGHV3-21, IGHV3-23, IGHV3-53, and IGHV3-73 while R2C2 could. Importantly, all alleles detected by R2C2 were also detected by TMI-seq data.

This analysis shows that, although specialized AIRR-seq protocols have an edge when detecting V segments that are rarely recombined, it produces incomplete and therefore often ambiguous V segment sequences. Although the TMI-seq method we developed can produce full-length V segment data, it requires a complex workflow and is therefore unlikely to be employed for routine clinical analysis. Overall, R2C2 presents an appealing set of tradeoffs and is therefore a promising tool for determining the V segment composition and usage within a sample.

#### Somatic Hypermutation

Fourth, we determined whether R2C2 reads would be accurate enough to detect the mutations in IGH sequences introduced by somatic hypermutation. We did this by comparing mutations in BCR transcript sequences which can undergo somatic hypermutation with TCR transcript sequences which are expected to be entirely free of somatic mutations.

We focused this analysis on mismatches which are by far the most common result of somatic hypermutation. We found the R2C2 reads did show only about two mismatches per 300nt TCR which corresponds to a mismatch-rate of 0.6% and is in line with the remaining 2% total error-rate in R2C2 reads being mostly composed of indels. Two mismatches per V segment can therefore be seen as background error in the potential mutated BCR sequences we analyzed next. IGHM transcripts are thought to be mostly expressed by naive unmutated B cells but can be also undergo somatic hypermutation. Here, we took advantage of the ability of R2C2 reads to distinguish membrane-bound and secreted (antibodies) isoforms of BCR transcripts. Secreted IGHM sequences contained more mutations than membrane-bound IGHM sequences, indicating that they are more likely to be expressed by B cells that have undergone activation and somatic hypermutation (Fig. 5A). This difference in mismatch levels disappears in IGHA1 BCR sequences which are known to be expressed only by B cells that have undergone activation and somatic hypermutation.

**Fig. 5:**
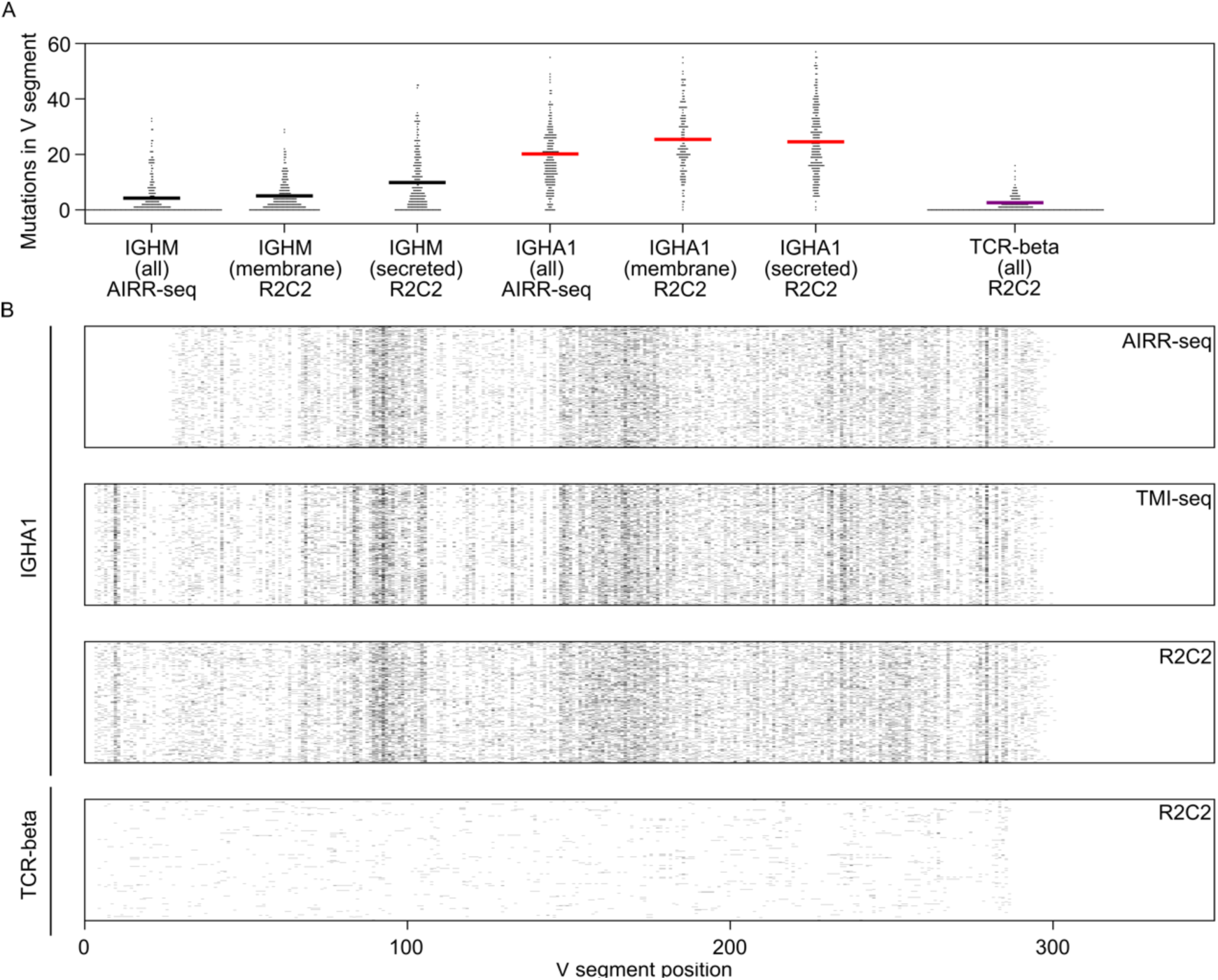
Somatic Hypermutation can be characterized within R2C2-based IGH repertoires. A) Mismatch mutations per IGH sequence as determined by IgBlast are shown as swarmplots separated by Isotype (IGHM, IGHA1), isoform (membrane-bound, secreted), and technology (R2C2, AIRR-seq) and compared to TCR-beta sequences. Averages are indicated by colored lines. B) The pattern of mutation locations in V segments in AIRR-seq, TMI-seq and R2C2 sequences is shown for 1000 randomly sampled IGHA1 or TCR-beta sequences.

Interestingly, mismatch levels were significantly higher in R2C2-derived IGHA1 BCR sequences than in AIRR-seq derived IGHA1 BCR sequences (Average 24.89 to 20.19, Monte-Carlo Permutation test p- value<0.00001). Adding randomly sampled R2C2-specific background-level mismatches observed in TCR-beta sequences as well as mismatches observed in the first 20 bases of R2C2 IGHA1 BCR sequences to AIRR-seq sequences does not abolish this significant difference (Average 24.89 to 23.8, Monte-Carlo Permutation test p- value<0.00001). One possible explanation of this may be that the primers employed by AIRR-seq fail at binding highly mutated V segments which then are not amplified and detected. In turn, this would indicate that R2C2 might have an advantage when investigating highly mutated sequences like those involved in the immune response to HIV^43^.

Finally, like AIRR-seq and TMI-seq, IGHA1 transcript sequenced by R2C2 show the mutational pattern characteristic of somatic hypermutation with mutational hotspots in CDR1 and CDR2. In contrast to AIRR-seq which uses amplicons primed from FR1 in the BCR transcript, TMI-seq and R2C2 sequence detect mutations all the way to the beginning of the V segment (Fig. 5B).

#### Clonality

Fifth, we investigated the ability of R2C2-based repertoires to capture the clonal composition of B cells in a sample. The BCR IGH sequences in a sample can be organized into clones (or lineages), i.e. sequences that are expressed by B cells belonging to the same B cell clone. B cell clones originate from a single naive B cell that is activated and starts proliferating. This proliferation is most often associated with somatic hypermutation and class-switching. Big lineages are therefore likely to be composed of class-switched sequences with similar but not identical mutation patterns. Most importantly, they have highly similar CDR3 sequences which can be used to group IGH sequences into lineages computationally. We performed this analysis for AIRR-seq and R2C2-based repertoires.

The lineages in the two repertoires were closely related (Fig. 6A) with 178 of the top 200 lineages in the AIRR-seq repertoire also being present in the R2C2-based repertoire. As expected, most of these repertoires had class-switched to the IGHA1 isotype and many contained additional lineage-specific mutations.

**Fig. 6:**
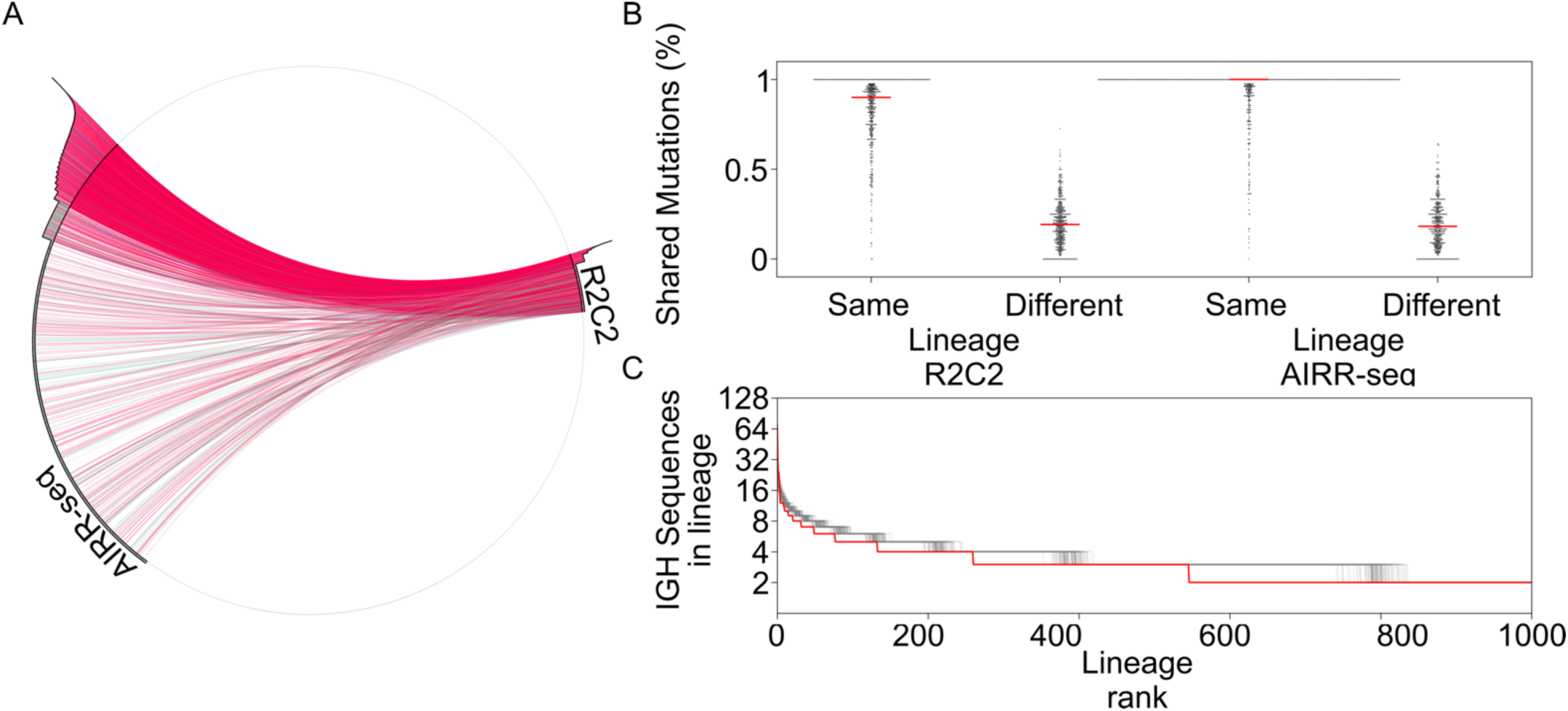
Clonal lineages can be measured by R2C2-based repertoires. A) Lineages shared between AIRR-seq and R2C2-based repertoires are indicated by connections within this Circos plot. Abundance of each lineage within each repertoire is shown as a histogram on the outside of the circle. B) The distribution of the percentage of mutations in an IGH sequence being shared with IGH sequences within the same or different lineages is shown as swarm-plots for R2C2-based and AIRR-seq repertoires. C) The size of lineages ordered by rank is shown for the R2C2-based repertoire (red) and 100 repertoires subsampled from the AIRR-seq repertoire to match the R2C2-based repertoires depth (grey)

Next, we used these mutations to confirm that IGH sequences were not spuriously grouped into lineages. To this end, we determined the percentage of mutations in each IGH sequence that are shared with other sequences within the same lineage or sequences in different sequences. For both R2C2 and AIRR-seq sequences, this percentage was much higher when comparing sequences within the same lineage confirming the overall accuracy of our lineage grouping approach. Finally, to see whether AIRR-seq and R2C2-based repertoires behave similarly when grouped into lineages, we repeatedly subsampled the AIRR-seq repertoire to the depth of the R2C2-based repertoire before grouping the subsampled sequences into lineages (Fig. 6C). The resulting AIRR-seq lineages were slightly larger than R2C2-based lineages at the same depth indicating that the 2% residual sequencing error present in R2C2 reads causes some related sequences to not be grouped into lineages.

Overall, analysis of AIRR-seq and R2C2-based lineages shows high concordance between the two methods. Of clinical relevance, R2C2-based lineages should be more than capable of tracking B cell clones within and between samples to for example tracking minimal residual disease in leukemia.

In its entirety, the analysis of the ability of R2C2 to investigate BCR IGH repertoires shows that R2C2 can replace specialized AIRR-seq for many purposes. While it still suffers from Indel errors, it can be used to determine V segment compositions, isotype usage, and somatic hypermutation levels and patterns, as well as the clonal structure of a population of B cells. Moreover accuracy of primary nanopore sequencing data is improving at a tremendous pace suggesting that Indel and other errors that are currently an issue for this application will rapidly decrease. Finally, while our R2C2/AIRR-seq comparison was focused on BCR IGH as the most complex and challenging to analyze AIR repertoire, our results suggest that R2C2-based repertoires for the other adaptive immune receptor loci are likely to be of comparable quality.

## DISCUSSION

As genomic analysis of samples becomes an integral component in clinical care, minimizing the number of separate assays that have to be performed and maximizing the information extracted from those assays that are performed should be a top priority. Here, we showcase the power of our R2C2 full-length cDNA sequencing approach for the in-depth analysis of PMBCs isolated routinely from blood. R2C2 libraries and Smart-seq2 can be generated from the same <1ng of total RNA making it possible to use less sample for transcriptome analysis. Further we recently developed a method for the depletion of hemoglobin transcripts from whole blood transcriptomes which would allow this type of analysis to be economically performed on whole blood RNA collected using e.g. PAXgene tubes^23^. If implemented on the ONT MinION, as we have done here, R2C2 generates reads of full-length cDNA at 98% accuracy at a cost of about ∼$200 per 1 million reads. Generating ∼10 million reads as we have done here is therefore not only feasible but likely to become routine.

We show that whole transcriptome analysis by R2C2 can be used to replace or enhance several specialized assays for the in-depth analysis of the complex transcriptomes of human immune cells. First, R2C2 full-length cDNA data can be used to generate allele-specific isoform-level transcriptome analysis which is beyond the capabilities of short-read data and impacted by high error rates and incomplete reads in standard ONT as well as low throughput in PacBio IsoSeq approaches. Second, we introduce a workflow that can complement short-read RNA-seq data to generate accurate allele-resolved full-length isoforms including the isoforms of all major HLA genes. Beyond accurate high-resolution HLA-typing, this approach will be very powerful for the identification of new HLA alleles present in the human population as it does not rely on databases of known HLA alleles as short-read based HLA-typing methods do. Fourth and finally, we identified tens of thousands of full-length adaptive immune receptor transcripts in our R2C2 data that can be compiled into repertoires containing a plethora of valuable data about the state of the adaptive immune system. R2C2-based repertoires therefore represent a convenient alternative to specialized AIRR-seq assays for the generation of AIRR-data which has been used in conjunction with specialized software^41,44^ to detect minimal residual disease^5^ and organ rejection^12^, or for basic research to track B cell clonal lineages or analyze immune aging^21^ or class-switching^20^. In contrast to AIRR-seq methods, generation of R2C2-based repertoires requires no specific primer sets which makes it a powerful tool for the investigation of not only human but vertebrate adaptive immune receptor diversity.

In summary, R2C2 full-length cDNA is a promising approach for the in-depth analysis of the human immune system and has the potential to replace or enhance specialized RNA-seq, HLA-typing and AIRR-seq approaches in the analysis of clinical samples. We showcase this potential in this proof-of-concept study to provide a stepping-off point for future studies to leave behind the limitations that short-read RNA-seq imposed on data generation and analysis.

## METHODS

### Sample collection and preparation

All experiments were approved by the Internal Review Board at the University of California Santa Cruz. Two whole blood samples were collected from a healthy human adult volunteer by the University of California Santa Cruz Student Health Center approximately six months apart. Samples were processed by Ficoll gradient (GE Healthcare) to extract PBMCs which were stored in liquid Nitrogen. RNA was extracted from one PBMC sample using the RNeasy mini kit (Qiagen). DNA was extracted from the other PBMC sample using the MagAttract HMW DNA kit (Qiagen).

### HLA-typing

HLA-typing was performed at the Immunogenetics and Transplantation Laboratory at University of California, San Francisco.

#### Sample preparation

DNA was quantified with NanoDrop (ThermoFisher Scientific, CA, USA) and adjusted to a concentration of 30ng/μL. Quality of DNA was assessed by measuring absorbance at A_230_, A_260_ and A_280_. DNA samples were amplified by long-range PCR using the Omixon Holotype HLA genotyping kit, generating full-length gene amplicons for HLA-A, B, C, DRB1, DRB3, DRB4, DRB5, DQA1, DQB1, DPA1 and DPB1 loci. Following PCR, amplicons were cleaned with Exo-SAP (Affymetrix, Santa Clara, CA, USA), quantified with QuantiFluor dsDNA system (Promega, Madison, WI, USA), and normalized to approximately 70 ng/μL.

#### Library preparation and sequencing

Sequencing libraries were generated for each sample using the Omixon Holotype HLA Genotyping Kit (Omixon, Inc. Budapest, Hungary). In brief, libraries from individual HLA amplicons were prepared by enzymatic fragmentation, end repair, adenylation, and ligation of indexed adaptors. The indexed libraries were pooled and concentrated with Ampure XP beads (Beckman Coulter) prior to fragment size selection using a PippinPrep(tm) (Sage Science, Beverly, MA, USA), selecting a range of fragments between 650 and 1300 bp. The size-selected library pool was quantified by quantitative PCR (qPCR; Kapa Biosystems, Wilmington, MA, USA) and adjusted to 2 nmol/L. The library was then denatured with NaOH and diluted to a final concentration of 8 pmol/L for optimal cluster density and 600 μL was loaded into the MiSeq reagent cartridge (v2 500 cycle kit). The reagent cartridge and flow cell were placed on the Illumina MiSeq (Illumina, San Diego, CA, USA) for cluster generation and 2 × 250 bp paired-end sequencing. Samples were demultiplexed on the instrument and the resulting FASTQ files were used for further analysis. HLA genotyping was assigned using TwinTM version 2.0.1 (Omixon, Inc. Budapest, Hungary) and IMGT/HLA database version 3.24.0_2, using 16000 read-pairs.

### Transcriptome sequencing library preparation

Approximately 200ng of total RNA was used to generate full-length cDNA using a modified Smart-seq2 protocol^9^. In short, RNA was reverse transcribed using Smartscribe RT (Clontech) and ISPCR-OligodT primer ISPCR-TSO (Table S1). Remaining RNA and primer dimers were digested and cDNA was PCR amplified using RNAseA, Lambda Exonuclease (NEB) and Kapa Biosystems HiFi HotStart ReadyMix (2X) (KAPA) with the following heat-cycling protocol: 37°C for 30 minutes, 95°C for 30 seconds followed by 12 cycles of (98°C 20 seconds; 67°C 15 seconds; 72°C for 10 minutes). The reaction was then purified using SPRI beads at a 0.85:1 ratio and eluted in H_2_O. The resulting full-length cDNA was then used as input into Smart-seq2, R2C2, and Smar2C2 library preparation protocols.

#### Smart-seq2

Full-length cDNA was then tagmented with Tn5 enzyme^25^ custom loaded with Tn5ME-A/R and Tn5ME-B/R adapters. The Tn5 reaction was performed using 50ng of cDNA in 5ul, 1 µl of the loaded Tn5 enzyme, 10 µl of H_2_O and 4 µl of 5× TAPS-PEG buffer and incubated at 55°C for 5 min. The Tn5 reaction was then inactivated by the addition of 5 µl of 0.2% sodium dodecyl sulphate and 5 µl of the product was then nick-translated at 72°C for 6 min and further amplified using KAPA Hifi Polymerase (KAPA) using Nextera_Primer_A and Nextera_Primer_B (Table S1) with an incubation of 98°C for 30 s, followed by 13 cycles of (98°C for 10 s, 63°C for 30 s, 72°C for 2 min) with a final extension at 72°C for 5 min. The resulting Illumina library was size selected on an agarose gel to be within 200-400bp and sequenced on a Illumina Nextseq 2×150 run.

#### R2C2

##### Splint Generation

23ul of H2O, 25ul of Kapa Biosystems HiFi HotStart ReadyMix (2X) (KAPA), 1ul of UMI_Splint_Forward (100uM) and 1ul of UMI_Splint_Reverse (100uM) were incubated at 95°C for 3 minutes, 98°C for 1 minute, 62°C for 1 minute, and 72°C for 6 minutes). The DNA splint was then purified with the Select-A-Size DNA Clean and Concentrator kit (Zymo) with 85ul of 100% EtOH in 500ul of DNA binding buffer.

##### Circularization of cDNA

200ng of cDNA was mixed with 200 ng of DNA splint and 2x NEBuilder HiFi DNA Assembly Master Mix (NEB) was added at the appropriate volume. This mix was incubated at 50C for 60 minutes. To this reaction we added 5ul of NEBuffer 2, 3ul Exonuclease I, 3ul of Exonuclease III, and 3ul of Lambda Exonuclease (all NEB) and adjusted the volume to 50ul using H_2_O. This reaction was then incubated 37°C for 16hr followed by a heat inactivation step at 80°C for 20 minutes. Circularized DNA was then extracted using SPRI beads with a size cutoff to eliminate DNA <500 bp (0.85 beads:1 sample) and eluted in 40 μL of ultrapure H_2_O.

##### Rolling circle amplification

Circularized DNA was split into four aliquots of 10 μL, and each aliquot was amplified in its own 50-μL reaction containing Phi29 polymerase (NEB) and exonuclease resistant random hexamers (Thermo) [5 μL of 10× Phi29 Buffer, 2.5 μL of 10 uM (each) dNTPs, 2.5 μL random hexamers (10 uM), 10 μL of DNA, 29 μL ultrapure water, 1 μL of Phi29]. Reactions were incubated at 30 °C overnight. T7 Endonuclease was added to each reaction which were then incubated at 37°C for 2h with occasional agitation. The debranched DNA was then extracted using SPRI beads at a 0.5:1 ratio and eluted in 50 μL of H2O.

##### Oxford Nanopore Technologies Sequencing

The resulting DNA was sequenced across four separate ONT MinION 9.4.1 flow cells. For each run, 1ug of DNA was prepared using the LSK-109 kit according to the manufacturer’s instructions with only minor modifications. End-repair and A-tailing steps were both extend from 5 minutes to 30 minutes. The final ligation step was also extended to 30 minutes. Each run took 48 hours and the resulting data in Fast5 format was basecalled using the high accuracy model of the gpu accelerated Guppy algorithm (version 2.3.5+53a111f, config file: dna_r9.4.1_450bps_flipflop.cfg). To generate R2C2 consensus reads, the resulting raw reads were processed using our C3POa pipeline (https://github.com/rvolden/C3POa).

#### Smar2C2

Library prep for this protocol is highly similar to Smart-seq2, however instead of cDNA, it uses the debranched rolling circle amplified DNA that is composed of cDNA concatemers. 50ng of this DNA was tagmented with Tn5 enzyme^25^ custom loaded with Tn5ME-A/R and Tn5ME-B/R adapters. The Tn5 reaction was performed using 50ng of cDNA in 5uL, 1 µL of the loaded Tn5 enzyme, 10 µl of H_2_O and 4 µl of 5× TAPS-PEG buffer and incubated at 55°C for 5 min. The Tn5 reaction was then inactivated by the addition of 5 µl of 0.2% sodium dodecyl sulphate and 5 µl of the product was then nick-translated at 72°C for 6 min and further amplified using KAPA Hifi Polymerase (KAPA) using Nextera_Primer_A and Nextera_Primer_B (Table S1) with an incubation of 98°C for 30 s, followed by 13 cycles of (98°C for 10 s, 63°C for 30 s, 72°C for 2 min) with a final extension at 72°C for 5 min. The resulting Illumina library was size selected on an agarose gel to be within 200-400bp and sequenced on an Illumina Nextseq 2×150 run.

#### AIRR-seq

200ng of total RNA was used for cDNA Smartscribe (Clontech) first-strand synthesis using a primer pool specific to the first exon of all IGH isotypes (IGHM, IGHD, IGHG1-4, IgGHA1-2, IGHE; see Table S1). In a two-cycle PCR reaction, second and third cDNA strands were synthesized using Kapa Biosystems HiFi HotStart ReadyMix (2X) and two modified primer pools complementary to the beginning of the Framing region 1 (FR1) of the V segment and ∼100 bp into the first exon of all IGH isotypes. All primers used in this two-cycle PCR reaction were modified to have unique molecular identifiers and partial Nextera sequences on their 5’ end. cDNA was purified and size-selected to >300nt using Select-A-Size DNA Clean and Concentrator kits (Zymo). In a 20-cycle PCR reaction, the cDNA is then amplified with primer completing Nextera sequences as well as Illumina i5 and i7 indexes to enable multiplexing of the libraries. Libraries were then sequenced on the Illumina MiSeq using a 2×300 run.

### Data Analysis

#### Gene Expression

R2C2 reads were aligned to the hg38 version of the human genome using minimap2^29^ using “-ax splice” and “-- secondary=no” flags and other standard settings. Smart-seq2 and Smar2C2 reads were aligned to the same genome sequence using STAR (version 2.7.1a)^45^ and an index built using the gencode v29 annotation gtf file. Read alignments were converted in gene expression counts using featureCounts^30^.

#### Isoform Analysis

R2C2 reads were analyzed to identify isoforms using version 3 of Mandalorion^9,10^ and standard settings. Isoforms were categorized using the sqanti_qc.py script of the SQANTI^31^ program with slight modifications to make it compatible with Python3. Isoform features were extracted from the categorized isoforms using custom scripts.

#### Allele-specific Isoforms

SNP present in the sample donor’s genome were identified using RNA-seq (Smart-seq2+Smar2C2) read alignments and GATK (version 3.8-1-0-gf15c1c3ef) following the standard RNA-seq SNP identification workflow (https://software.broadinstitute.org/gatk/documentation/article.php?id=3891). Homozygous SNPs were discarded. The remaining heterozygous SNP were phased using R2C2 reads and the new Mandalorion utility TurboPhaser.py, taking advantage of R2C2 reads spanning entire gene loci and grouping SNPs that appeared in the same reads. TurboPhaser.py also sorted R2C2 reads and RNA-seq reads into alleles based on the SNPs they contained. The sorted R2C2 reads were then used in the Mandalorion pipeline to identify isoform sequences. The sequences were then error-corrected using pilon^*46*^ and RNA-seq reads aligned to the isoform sequences using minimap2 with the “-x sr” preset.

#### HLA-typing

Using the HLAtyping.py script Allele-specific error-corrected isoform sequences were aligned to the human genome using minimap2 (“--secondary=no -x splice”). Isoforms aligning to HLA loci were then realigned to HLA transcript sequences retrieved from the IPD-IMGT/HLA database^37,38^ using minimap2 (“-ax splice -N 100”). For each HLA gene and allele, the best full-length match was reported. For RNA-seq-only approaches, Smart-seq2 reads and Smar2C2 reads were pooled and processed as required by seq2HLA and arcasHLA and the programs were run with standard settings.

#### AIRR analysis

R2C2 reads aligning to adaptive immune receptor loci were extracted using samtools^47^. The sequences were then analyzed using *IgBlast*^*39*^ with output format 19 with V, D, and J segments retrieved from IMGT^48^. For the in-depth analysis of BCR IGH repertoires this output was then parsed using custom scripts to report CDR3 length, Vsegment, isotype, and isoform (secreted vs membrane-bound). Sequences were then grouped into lineages using a simple single linkage clustering approach using a 90% CDR3 nucleotide similarity cut-off.

## Data and Code Availability

All sequencing data generated for this study is available at SRA (PRJNA559668). TMI-seq sequences are available at SRA (PRJNA291102). Processed R2C2 and AIRR consensus reads are also available at https://users.soe.ucsc.edu/~vollmers/PBMC_data/R2C2_reads.fa and https://users.soe.ucsc.edu/~vollmers/PBMC_data/AIRRseq_reads.fasta Mandalorion and its utilities for isoform identification and sequence determination are available on github (https://github.com/rvolden/Mandalorion-Episode-III). The Mandalorion github also contains scripts for processing SQUANTI classification, sorting R2C2 reads into alleles, and HLA-typing. C3POa generates R2C2 consensus reads from ONT raw reads and is available on github (https://github.com/rvolden/C3POa). The C3POa github also contains scripts for identifying and merging reads with matching UMIs. Scripts for the parsing and grouping AIRR data into lineages is available at (https://github.com/christopher-vollmers/AIRR).

## Data Visualization

All data visualization was done using *Python*/*Numpy*/*Scipy*/*Matplotlib*^*49–52*^. Schematics were drawn in Inkscape (https://inkscape.org/en/)

## ACKNOWLEDGEMENTS

We thank Dr. Raja Rajalingam and the Immunogenetics and Transplantation Laboratory (ITL) at the University of California, San Francisco for performing high-resolution HLA-typing. We thank Ed Malloy, Aisha Coons, and Jerrold Michaud of the Student Health Center at the University of California Santa Cruz for expert assistance with blood draws. We acknowledge funding by the National Human Genome Research Institute/National Institute of Health Training Grant 1T32HG008345-01 (to C.C., A.B., and R.V.), the Hellman Foundation, Santa Cruz Cancer Benefit Group, and National Institute of General Medical Sciences/National Institute of Health Grant 1R35GM133569-01 (to C.V.)

## Author Contributions

C.C., A.B, and M.A. performed experiments. C.C., R.V., and C.V. analyzed the data. C.C. and C.V. conceived of the study and designed experiments. C.V., C.C., A.B., R.V., and M.A. wrote and edited the manuscript.

## Supplemental Information

**Fig. S1:**
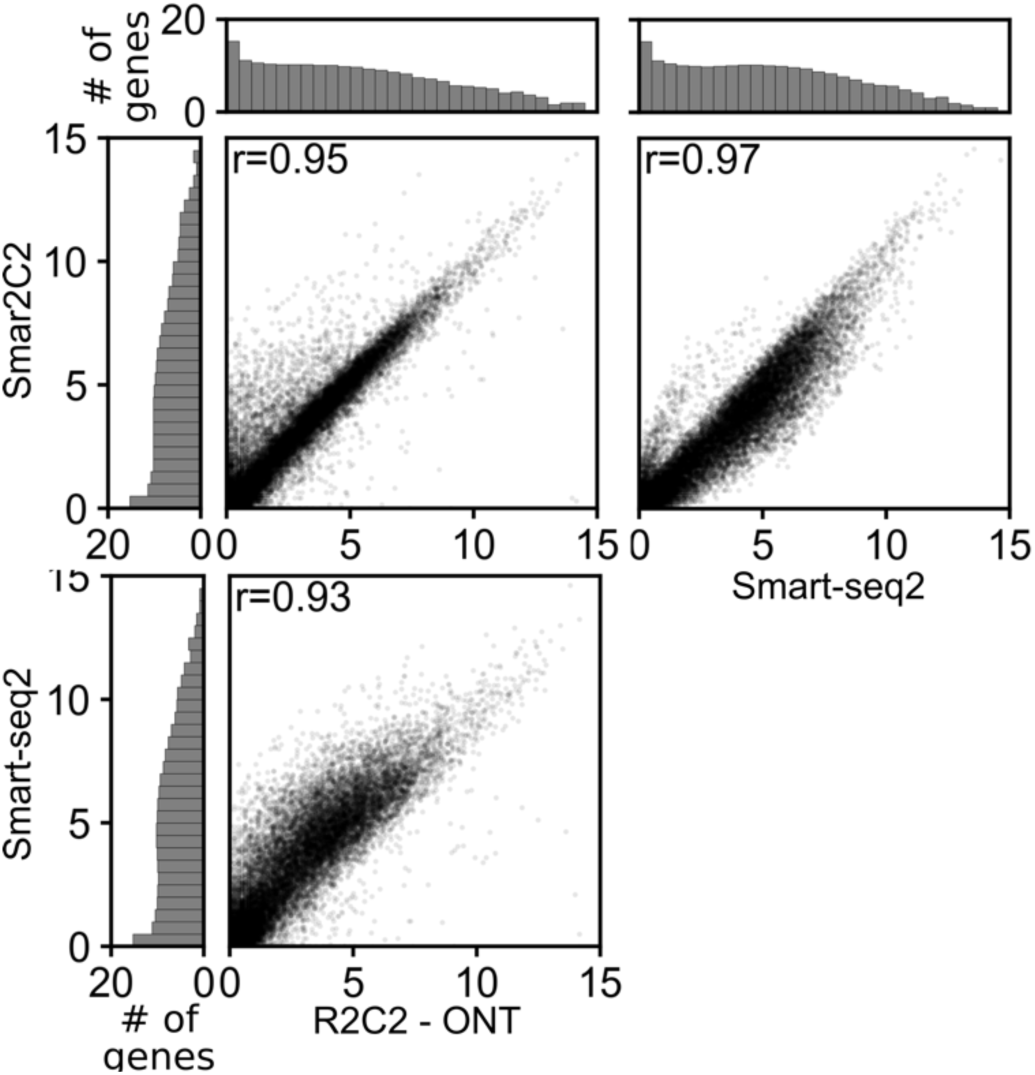
Gene Expression Quantification with different short- and long-read methods. We quantified gene expression using featureCounts and plotted the resulting data as scatter plots comparing R2C2, Smar2C2, and Smart-seq2. Histograms on the borders indicate the number of genes in each expression bin for the respective technology. R-values given are Pearson r. Gene expression values (RPM) and gene numbers are all shown as log2(value+1).

**Table S1:**
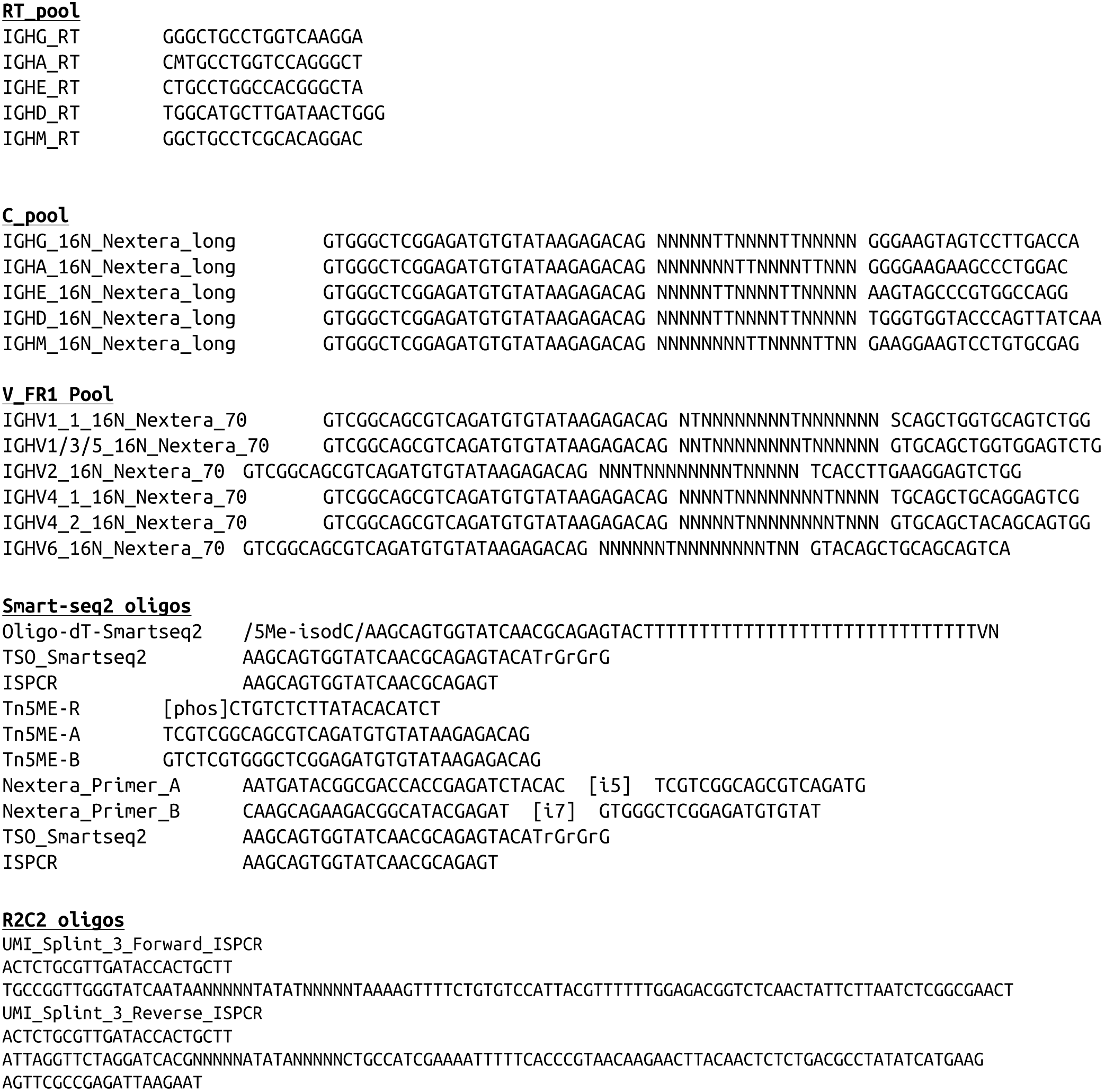
Oligonucleotide sequences used in this study.

**Table S2:**
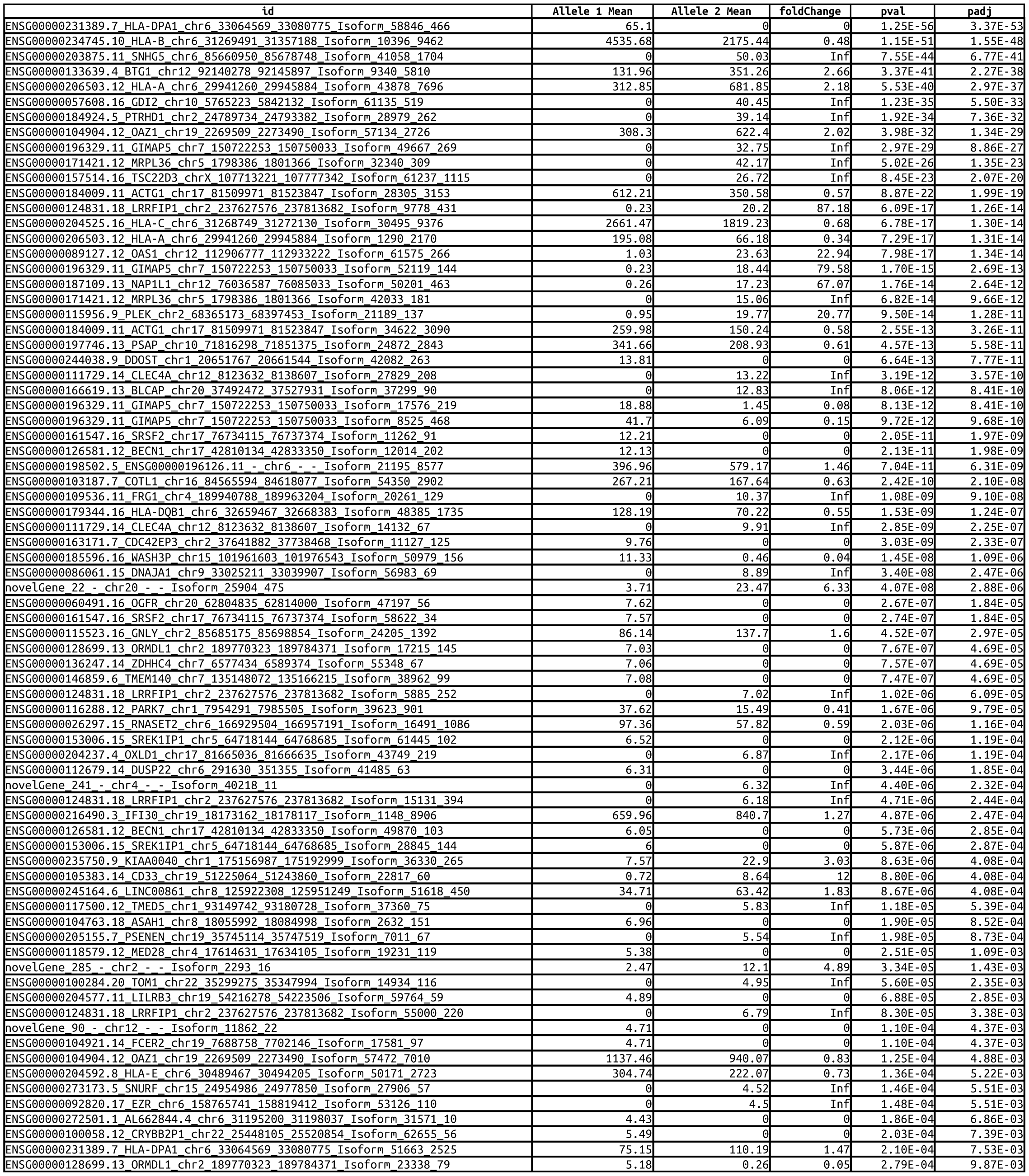
DESeq output evaluating differential isoform expression between alleles. Isoforms with an adjusted p-value <0.01 are shown.

